# Pyk2 suppresses contextual fear memory in a kinase-independent manner

**DOI:** 10.1101/216770

**Authors:** Lun Suo, Jin Zheng, Yuxiao Zhou, Liling Jia, Yanping Kuang, Qiang Wu

**Affiliations:** Center for Comparative Biomedicine, MOE Key Laboratory of Systems Biomedicine, Institute of Systems Biomedicine, SCSB, Shanghai Jiao Tong University, Shanghai, China; Department of Assisted Reproduction, Shanghai Ninth People’s Hospital, Shanghai Jiao Tong University School of Medicine, Shanghai, China; School of Life Sciences and Biotechnology, Shanghai Jiao Tong University, Shanghai, China

**Author notes:** These authors contributed equally to this work (Lun Suo, Jin Zheng, Yuxiao Zhou). (QW); (LS).

## Abstract

Post-traumatic stress disorder (PTSD) is a psychological illness characterized by recalling a feeling of distress when re-experiencing the original trauma-related cues. This associative fear response plays an important role in some psychiatric disorders and elucidation of the underlying mechanisms is of great importance. Here, we constructed *Pyk2* null mice and found that these mutant mice showed enhancement in contextual-fear memory, but no changes in auditory-cued and spatial-referenced learning and memory. Moreover, using kinase mutant mice, we observed that Pyk2 suppressed contextual fear memory in a kinase-independent pathway. Using high-throughput RNA sequencing, we found that immediate early genes (IEGs), such as *Npas4*, c*Fos*, *Zif268/Egr1*, *Arc*, and *Nr4a1*, were enhanced in *Pyk2* null mice. We further demonstrated that Pyk2 disruption affected pyramidal neuronal complexity and spine dynamics. Thus, we demonstrated that Pyk2 is a novel fear memory suppressor molecule and *Pyk2* null mice provides a model for understanding fear-related disorders.

## Introduction

Fear responses to environmental threats endow human beings to seek comfort and to avoid danger, thus increasing the chances for survival. Dysfunction in fear memory maintenance or retrieval has evolutionary disadvantages, whereas hyperfunction in this capacity leads to mental illnesses, including PTSD or phobia responses. Thus, it is required for human beings to maintain a dynamic balance between fear-memory strengthen and suppression. For memory strengthen, a number of molecules including neurotransmitters [1], transcription factors [2], and intracellular signaling molecules such as MAPK were found to be important [3]. In contrast, multiple repressive molecules and signaling pathways are important in fear memory suppression [4].

For the assembly of neural circuits in the brain, the clustered protocadherins (Pcdhs)-cell adhesion kinases (CAKs) signaling pathway might play critical roles in keeping a balance between neurite connection and repulsion. In particular, two cell-adhesion kinases, Pyk2 (also known as CAKβ or PTK2B) and Fak, function downstream of the clustered Pcdh proteins to regulate cytoskeletal reorganization [5–7]. Pcdhs are a large family of diverse cell-adhesion proteins that are specifically expressed on the cell surface of neurons in the brain [8,9]. Individual neurons only express a particular subset of clustered *Pcdh* genes determined by specific long-distance promoter-enhancer looping interactions within a CTCF/cohesin-mediated topological chromatin domain [10,11]. Homophilic adhesion between extracellular ectodomains of clustered Pcdh proteins are thought to be essential for proper assembly of neuronal connectivity [12,13]. For example, homophilic recognition between the same sets of clustered Pcdh proteins may lead to repulsion, thus self-avoidance and even spacing, between neurites from same neurons through cytoskeletal rearrangement [14,15]. The mechanisms underlying the adhesive recognition-lead repulsion is not known but may involve signaling molecules such as cell-adhesion kinases.

Pyk2 is a non-receptor tyrosine kinase and is very abundantly expressed in the hippocampus of postnatal rats, reaching maximal levels in adulthood [16]. We and others recently found that Pyk2 plays important roles in dendritic and spine morphogenesis [7,17,18]. Pyk2 is post-synaptic located and interacts with NMDA receptors [19] or PSD95 [20], implicating a role of Pyk2 in both LTP [21,22] and LTD [23]. Recently, Pyk2 was also identified as a susceptibility locus for Alzheimer diseases and dementias based on genome-wide association studies [24,25]. Here, we found Pyk2 as a repressor of contextual fear memory based on behavior studies using gene modified mice by CRISPR.

## Results

### Generation of *Pyk2* mutant mice and exclusive expression of Pyk2 in the hippocampus

To determine the physiological function of *Pyk2 in vivo*, we generated *Pyk2* mutant mice using CRISPR/Cas9 system. We designed sgRNA target sequence (gene ID: 19229 from 63961711 to 63961730) based on the first exon of the *Pyk2* locus (Fig 1A) and transcribed this sgRNA and Cas9 mRNA *in vitro*. After injection into fertilized eggs, two lines of frame-shifted mutant mice (*Pyk2* KO^-8bp^ and *Pyk2* KO^+1bp^) were identified. Germline transmission of the null allele was initially screened by BslI enzyme digestion of the PCR products from the tail DNA of pups (Fig 1B-1D), and then confirmed by Sanger sequencing.

**Fig. 1.**
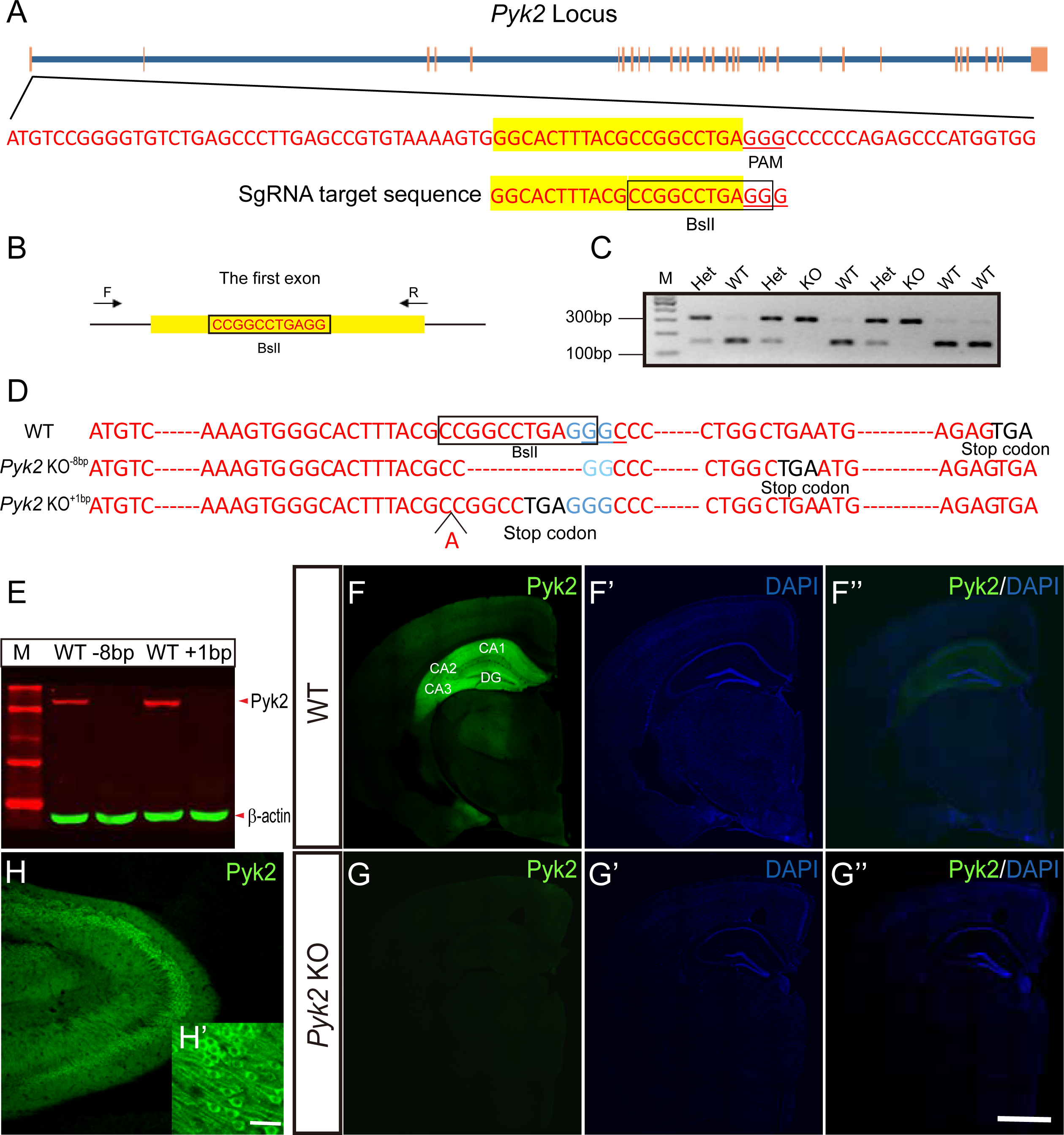
*Pyk2* gene disruption and its expression patterns in the brain of mice. (A) Schematic of the Cas9/sgRNA-targeting sequence of the *Pyk2* gene. The sgRNA-argeting sequence is highlighted in yellow and the protospacer-adjacent motif (PAM) is underlined. (B) The forward (F) and reverse (R) primers for PCR and the BslI restriction enzyme site at the target regions are marked. Electroporation of PCR products after estriction enzymes digestion (C) and confirmation by Sanger sequencing (D). (E) The validation of gene disruption was performed using Western blot in both *Pyk2* KO^-8bp^ and *Pyk2* KO^+1bp^. (F and G) Confocal images of brain sections from WT and *Pyk2*-KO mice mmunostained with antibodies against Pyk2 (F, G), DAPI (F’, G’) and merged (F”, G”). Bar in G” indicates 1,000 µm for F and G. CA1, 2, 3: cornu ammonis 1, 2, 3; DG: dentate gyrus. (H and H’) Enlarged pictures of hippocampus were indicated for pyramidal cells. Bar in H’ indicates 50 µm for H’ and 100 µm for H.

To validate the KO phenotype of mutant mice, we detected an immunoreactive band at 116 kDa corresponding to Pyk2 in hippocampal samples from wild-type, but not *Pyk2*-KO mice (Fig 1E). We also found that Pyk2 proteins were predominantly expressed throughout the adult hippocampus (Fig 1F), but not in embryonic 18.5 and P0 hippocampus (S1 Fig). In CA1, CA2, CA3, and DG of the hippocampus, Pyk2 was mainly found in the cytoplasm and dendrite of pyramidal neurons (Fig 1H and 1H’). In contrast, *Pyk2*-KO mice showed no signal (Fig 1G).

### Pyk2 suppresses contextual fear memory in a kinase-independent pathway

Hippocampus is an important organ for learning and memory in the central nervous system. We assessed two hippocampal-dependent learning and memory behavior tests, including associate fear conditioning [26] and Morris water maze [27,28] using wild-type and *Pyk2*-KO mice. For fear conditioning, mice were put into the training chamber for adaptive training on Day 1. Fear training was performed on Day 2 (Fig 2A). On Day 3, mice were put back into the training chamber to test contextual fear conditioning. This was followed by placing the animals into a novel chamber to test novel contextual fear conditioning or into a novel chamber and exposed to the original auditory cues to test trace fear conditioning. After repeated fear training using auditory cues on Day 2, we found increases in frequencies of fear responses but no differences between wild-type and *Pyk2*-KO mice (Fig 2B), indicating that memory acquisition and fear responses were intact in *Pyk2*-KO mice. On Day 3, fear response of *Pyk2*-KO mice was significantly increased as compared with their wild-type littermates when the mice were put back into the training context 24h after training (Fig 2C). In contrast, both *Pyk2*-KO and wild-type mice showed but without any phenotypic difference in freezing behavior when the animals were put into a novel context only (Fig 2C) or novel context together with the auditory tone (Fig 2D), indicating that *Pyk2*-KO mice might be more capable of contextual associative fear memory maintenance but not the auditory cued memory maintenance.

**Fig 2.**
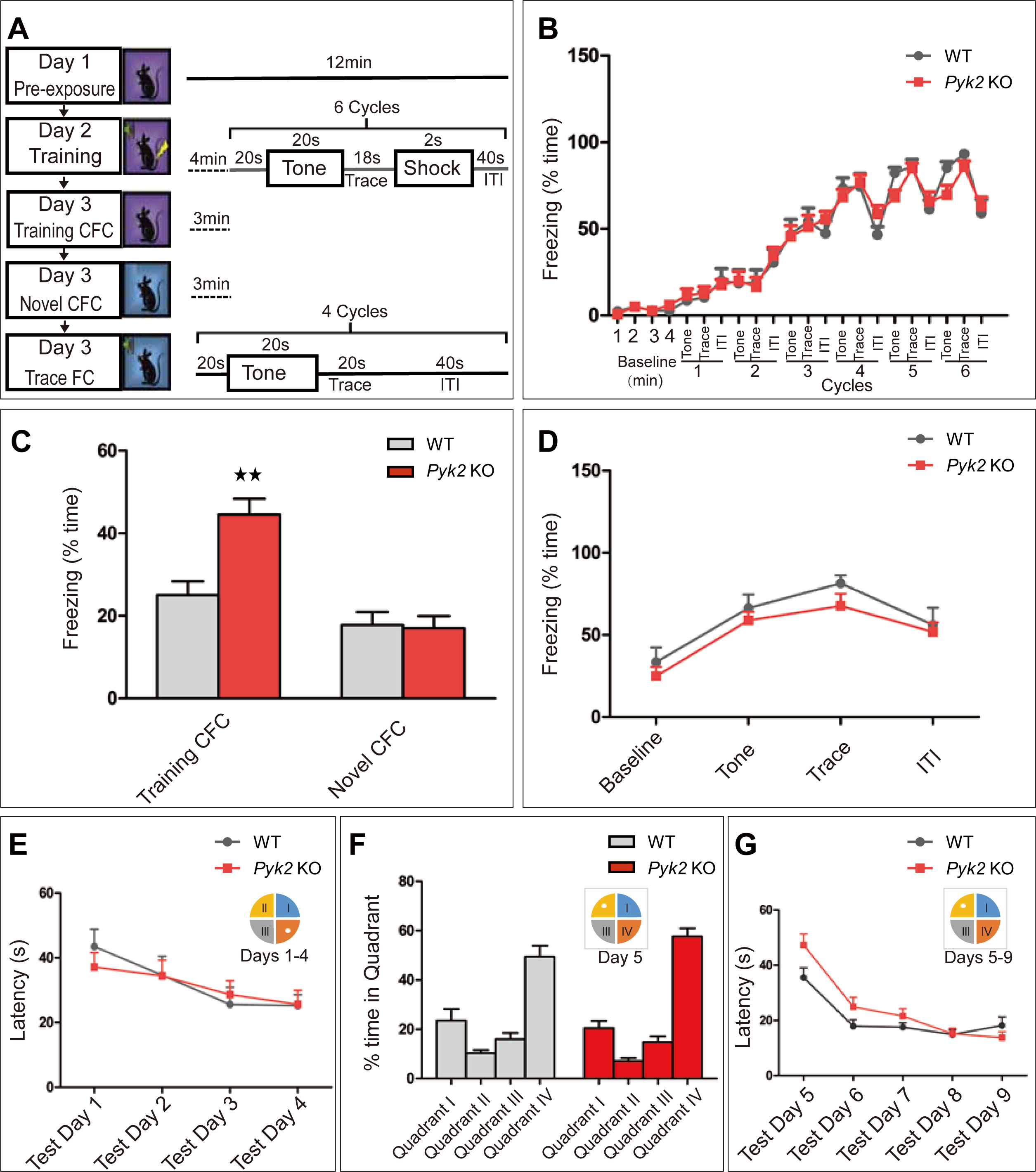
Hippocampal-dependent contextual fear memory was enhanced in *Pyk2*-KO mice. (A) The procedure for fear conditioning test. (B) Trace fear response in *Pyk2*-KO (n = 20) and WT (n = 12) mice were significantly enhanced during training but without significant difference between genotypes (F_times_(21,630) = 90.16, p < 0.0001; F_genotype_(1,630) = 0.01, p = 0.91). (C) *Pyk2*-KO mice exhibited significantly higher in fear response than that of WT mice (*Pyk2* KO = 44.55%, WT = 25%, p < 0.01) in training context 24h post training. n novel context, fear response showed no difference between genotypes (WT = 17.8% vs *Pyk2* KO = 17%, p > 0.05) (**p < 0.01). (D) Auditory trace fear conditioning was unaffected in *Pyk2*-KO mice 24h after training (F_sessions_(3,75) = 20.28, p < 0.0001; F_genotype_(1,75) = 1.45, p = 0.24; *Pyk2* KO, n = 17 and WT, n = 10). (E) During initial acquisition of a hidden platform location, escape latencies were similar between *Pyk2*-KO (n = 22) and wild-type mice (n = 13) (F_times_(3,99) = 7.58, p = 0.0001; F_genotype_(1,99) = 0.02, p = 0.89). (F) Both genotypes showed a significant preference for the target quadrant (quadrant IV) at test Day 5 (F_quadrants_(3,132) = 84.70, p < 0.0001). There was no significant difference between genotypes (F_genotype_(1,132) = 0.01, p = 0.92). (G) Following he reversal of platform location, there was also no significant difference between genotypes in escape latencies between test Days 5-9 (F_times_(4,132) = 40.45, p < 0.0001; F_genotype_(1,132) = 1.46, p = 0.24). Data are represented as mean µ SEM.

We also performed Morris water maze tests, consisting of 2 days training, 4 days of acquisition, followed by 5 days of memory testing. During initial spatial memory learning, we found that *Pyk2*-KO and wild-type mice exhibited similar escape latency durations during testing from Days 1 to 4 when the platform was put in quadrant IV (Fig 2E), indicating that the mice acquired spatial memory equally well between genotypes. When the hidden platform was moved to quadrant II of the pool on Day 5 of testing, both wild-type and *Pyk2*-KO mice showed a persistence of entries to the old platform (Fig 2F) and no difference was found between genotypes in escape latency times during test Days 5-9 (Fig 2G). These data indicated similar spatial memory formation and erasing between wild-type and *Pyk2*-KO mice.

Pyk2 is a tyrosine kinase and could transduce signalling through its catalytic activity mediated by tyrosine residue at 402. To this end, we generated a Pyk2 kinase site mutant mice with tyrosine residue 402 changed into phenylalanine (Pyk2^Y402F^) (S2A-S2C Fig). This kinase site mutant mice were then tested for contextual associated memory, auditory cued memory, and Morris water maize. Of interest, these behavior tests indicated that Pyk2^Y402F^ mice showed no difference in either fear memory acquisition (S2D Fig) or contextual fear memory (S2E Fig) and auditory cued fear memory(S2F Fig), as well as spatial cued memory (S2G-S2I Fig), indicating that Pyk2-controled contextual fear memory maintenance did not rely on its catalytic activity. Another critical residue of Pyk2, tyrosine residue 881, could couple Pyk2 to Grb2/SOS complex and activate specific mitogen-activated protein kinase (MAPK)/extracellular signal-regulated kinases (ERK) signalling pathway which has been reported to play critical regulatory roles in long-term potentiation dependent gene expression [29,30]

### Pyk2 represses immediate early genes

To test potential roles of Pyk2 in hippocampal gene regulation, we compared the transcriptome profiles of hippocampus between wild-type and *Pyk2*-KO mice using high-throughput RNA sequencing. According to cutoff threshold of >2 fold changes or <0.5 fold changes and FDR<0.001, a total number of 52 RNA transcripts were identified: 17 downregulated and 35 upregulated in *Pyk2*-KO mice compared with that of wild-type mice (Fig 3A-3C and S1 Table). Among the upregulated genes, we found neuronal activity regulated genes and early-response genes, including *Npas4*, c*Fos*, *Zif268/Egr1*, *Arc*, and *Nr4a1* (Fig 3C), which were illustrated in a representative UCSC browser image (Fig 3D). As previous studies reported, for normal fear conditioning training in mice, IEGs expression increased from 10 to 30 min and then followed by a significant decrease at 4 hours after training [4]. These gene expression patterns might be beneficent to keep a balance between memory consolidation and suppression. Interestingly, our work found that the expression of these genes were significantly increased when *Pyk2* was deleted (Fig 3E), indicating that *Pyk2* might act as a switch in regulating IEGs expression. Whereas, as expected, not any neuronal activity genes mentioned above were found to be differently expressed in the hippocampus between Pyk2^Y402F^ mutant and its wild-type littermate mice (S2 Table). That could explain why Pyk2^Y402F^ mutant mice show no enhancement in contextual-dependent fear memory maintenance.

**Fig 3.**
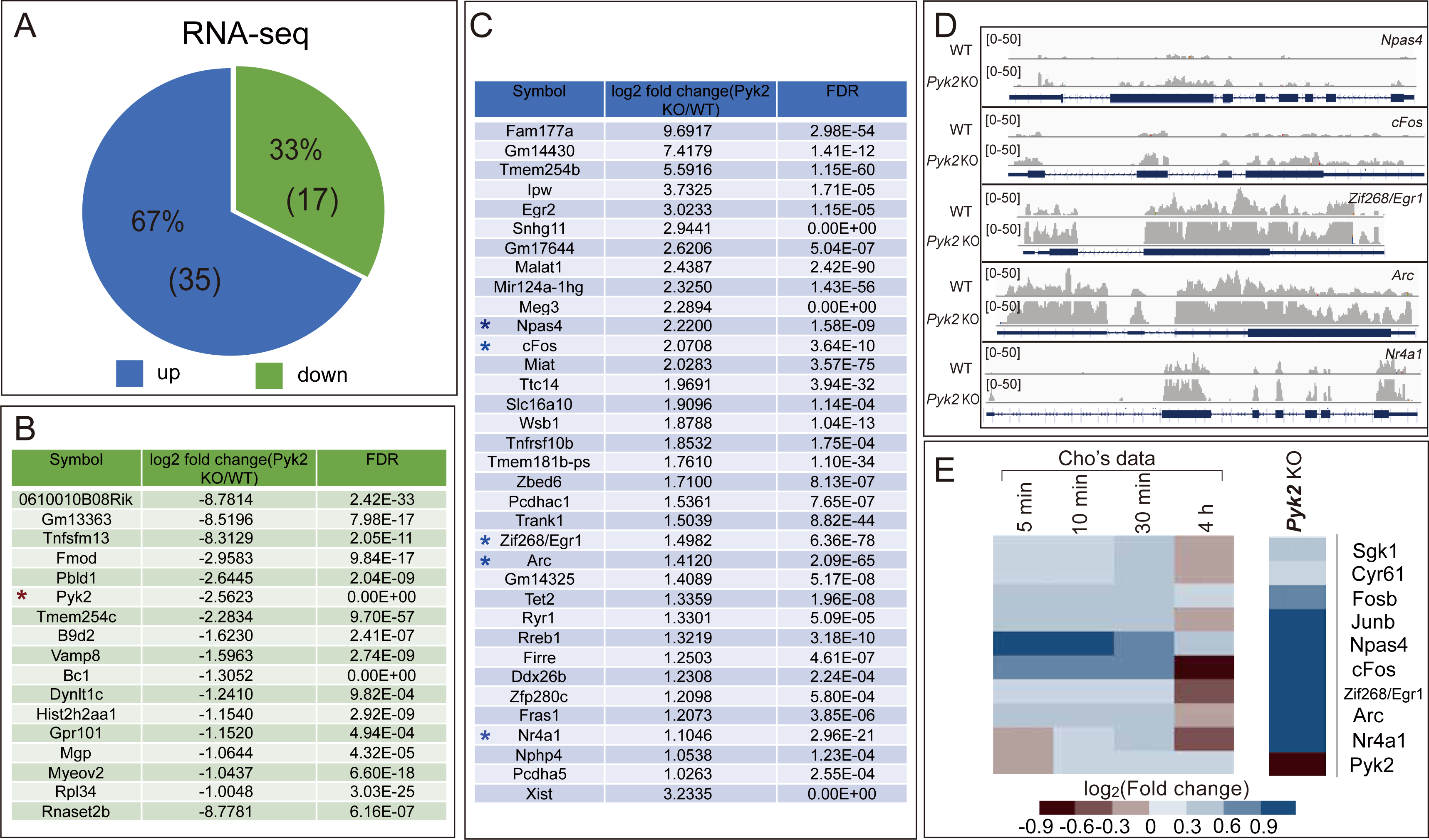
Loss of *Pyk2* activates expression of multiple neuronal activity related genes n the hippocampus. (A) Pie chart showing genes up- and down-regulated in hippocampus of *Pyk2*-KO mice. (B and C) 17 down-regulated (B) and 35 up-regulated (C) differentially-expressed genes n *Pyk2*-KO mice were ranked by fold change. *Pyk2* was marked with red asterisks and EGs were marked with blue asterisks. (D) UCSC genome browser images showing ncreases of transcription in *Pyk2*-KO mice by RNA-sequence of selected neuronal activity related genes. (E) Expression patterns of some IEGs in fear condition training under normal conditions (left panel from Cho’s sequencing data) or in *Pyk2*-KO mice (right panel from our sequence data).

### Morphological changes of hippocampal neurons in *Pyk2*-KO mice

To test whether disruption of Pyk2 affected neuronal morphology and synaptic plasticity, we performed Golgi staining of the pyramidal neurons from wild-type and *Pyk2*-KO littermate mice. As shown in Fig 4A-4D, hippocampal neurons in CA1, CA2, CA3, and DG regions from *Pyk2*-KO mice showed higher morphological complexity in dendrites than that of wild-type mice. Quantification of the reconstructed confocal images revealed significant increases in total length (Fig 4E), the number of branch points (bifurcations) (Fig 4F), and segments between branch points (Fig 4G) in *Pyk2-*KO mice.

**Fig 4.**
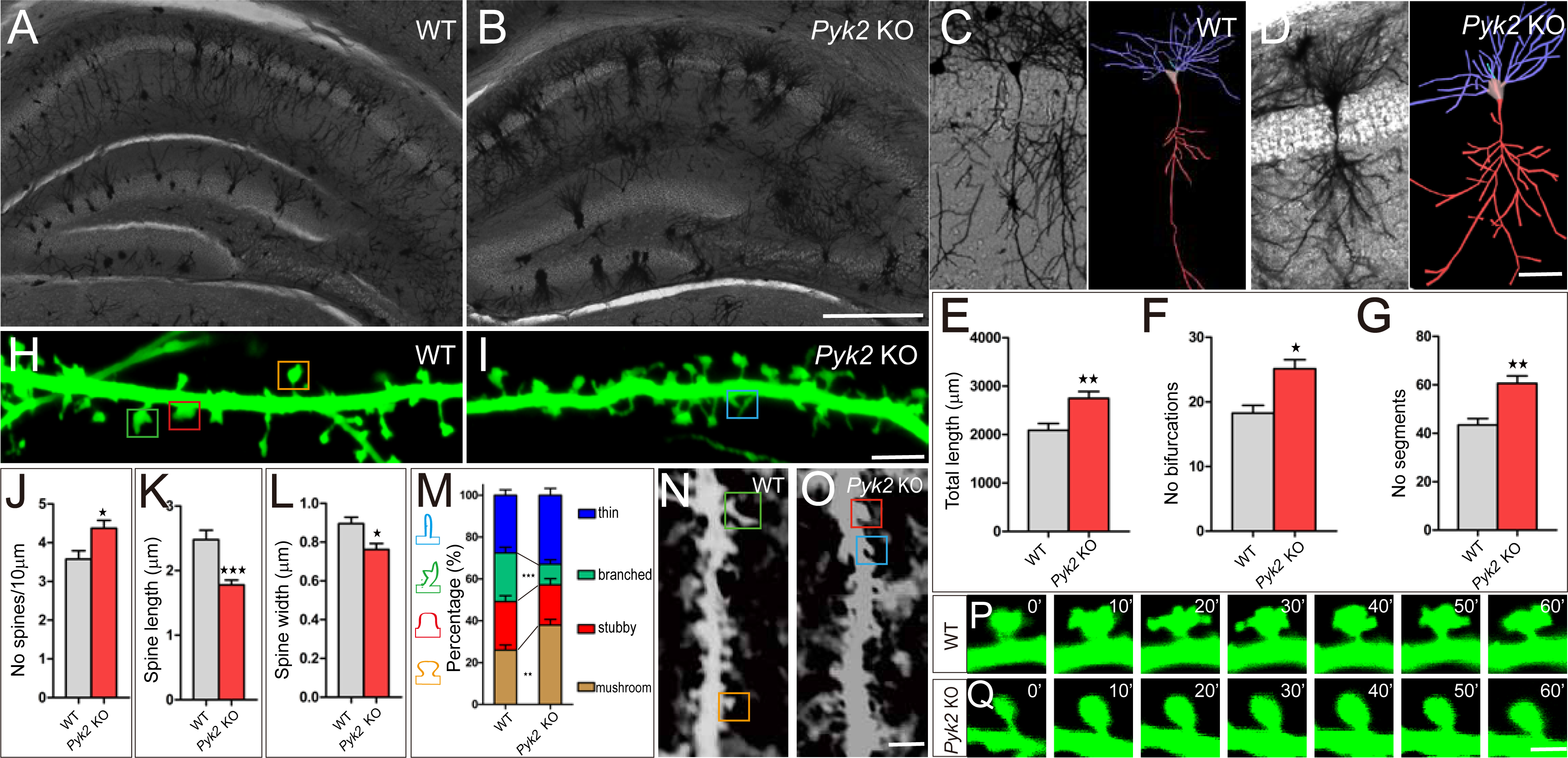
Morphology of hippocampal neurons in *Pyk2*-KO mice. (A-D) The representative Golgi staining sections and the reconstructed 3D confocal mages of CA1 hippocampal neurons from the WT (A, C) and *Pyk2*-KO (B, D) littermates. Bar in B represents 200 µm for A and B. Bar in D represents 50 µm for C and D. (E-G) Quantification of total length (E), branch points (bifurcation) (F) and the number of segments (G) between branch points (WT, n = 20; *Pyk2* KO, n = 27) (*p < 0.05, **p < 0.01). (H and I) Representative images of spines in dendritic segments of 14 DIV neurons rom WT (H) or *Pyk2*-KO mice (I). Bar in I represents 5 µm for H and I. (J-M) Quantification of the spine density (J), length (K), and width (L) (WT, n = 370 spines of 22 dendritic segments; *Pyk2* KO, n = 452 spines of 24 dendritic segments) and the percentages of spine morphology analysis (WT, n = 325 spines of 21 dendritic segments; *Pyk2* KO, n = 382 spines of 26 dendritic segments) (M) of 14 DIV neurons (*p < 0.05, **p < 0.01, ***p < 0.001). (N and O) Representative confocal images of the basal dendritic segments from he CA1 pyramidal neurons of the WT (N) and *Pyk2*-KO (O) mice. Bar in O represents 3 µm for N and O. (P and Q) Representative confocal images of spine structural dynamics of hippocampal neurons from WT (P) and *Pyk2*-KO (Q) mice in time-lapse experiments at every 10 min interval for a total of 1 hour. Bar in Q represents 2 µm for P and Q. Data are represented as mean ± SEM.

To further confirm the role of *Pyk2* in dendritic development, we transfected GFP-encoding plasmids into hippocampal neurons during primary cultur*e* for better tracing of dendrite morphology (Fig 4H and 4I). As shown in Fig 4J, the spine density of dendritic segments was significantly increased in *Pyk2*-KO mice compared with wild-type control mice. However, there were significant decreases in the spine length (Fig 4K) and spine width (Fig 4L) in *Pyk2*-KO mice. Furthermore, the proportion of spine with a classical head shape was significantly increased, whereas the spine with branched heads was significantly decreased in *Pyk2* mutants (Fig 4M-4O). Interestingly, time-lapse tests also indicated that the spine remodeling was more stable with less structural plasticity when *Pyk2* was deleted (Fig 4P and 4Q; S1 and S2 Movie), which could explain why *Pyk2* disruption induces contextual fear memory maintenance, as stable spine has been known as a potential structural basis for long-term memory storage [31]. Taken together, these data suggested a role of the Pyk2 protein in dendrite formation and spine dynamics, which was critical for fear memory.

## Discussion

The cellular and molecular mechanisms of fear memory acquisition and maintenance have been extensively studied in the amygdala [32,33], whereas some other brain structures including, superior colliculus [34], thalamus [35], and cortex [36,37], as well as hippocampus [38] also contribute to this biological process. Here, through analyzing genes expression, dendrite morphology, and synaptic structural plasticity, in conjunction with mouse genetics, we found Pyk2 as a contextual fear memory suppressor protein capable of regulating neuronal activity in a kinase-independent pathway.

Our *Pyk2*-KO mice grew up to the adult stage with normal body weight and showed predicted ratios of genotype in line with Mendelian inheritance, similar to the *Pyk2*-KO mice previously generated [18,39]. *Pyk2* is predominantly expressed throughout the CA1, CA2, CA3, and DG regions of the hippocampus in the adult mice (Fig 1) and rat [16] but not at the embryonic 18.5 and P0 stages, indicating that *Pyk2* is dispensable for embryonic neurogenesis, neuronal differentiation, and brain development. Based on the hippocampal-specific expression pattern, we speculate that Pyk2 might be a critical molecular component involved in cognitive behaviors and associated with psychiatric disorders relevant to dysfunction in learning and memory process.

Two classic hippocampal-dependent learning and memory tests, Pavlovian fear conditioning and Morris water maze, demonstrated that the acquisition of contextual-fear and spatial-reference memories is not affected in *Pyk2*-KO mice. By contrast, contextual fear memory maintenance was enhanced when *Pyk2*-KO mice were put back into the training context 24 hours after training, indicating that molecular mechanisms and neuronal circuitries responsible for contextual-cued fear memory is increased in *Pyk2*-KO mice, consistent with a suppressor role of PyK2 in memory maintenance. Pyk2, a close homologue of Fak, is a tyrosine kinase capable of regulating signaling pathways through its catalytic domain [40,41]. However, we did not find any difference in contextual-fear conditioning between Pyk2^Y402F^ kinase site mutant and wild-type control mice (Fig 2), suggesting that Pyk2 functions in fear memory suppression in a kinase-independent pathway.

Pyk2 has a known scaffolding function independent of its kinase activity through recruiting of the Grb2/SOS complex by the phosphorylated residue Y881 [29,40]. The Grb2/SOS complex regulates the MAPK/ERK-CREB signaling pathway [29]. Thus, Pyk2 might suppress contextual fear memory through regulating Grb2/SOS-MAPK/ERK-CREB signaling pathway, which plays regulatory roles in long-term potentiation dependent gene expression [30]. Through RNA-seq, we found that a number of CREB-regulated IEG genes, encoding neuronal-activity-related transcription factors NPAS4, cFOS, Zif268/ERG1, NR4A1, as well as activity-regulated cytoskeletal associated protein (ARC), were significantly increased in *Pyk2*-KO mice at transcriptional levels. Remarkably, none of these IEGs were found to be significantly increased in the Pyk2^Y402F^ mice, consisting with none behavioral phenotype in the Pyk2^Y402F^ mice. The IEGs have been widely implicated in hippocampal-dependent learning and memory and is believed to play an integral role in synapse-specific plasticity [42].

Our study indicated that neurons from *Pyk2*-KO mice showed more complexity during dendrite arborization. In addition, the proportion of neuronal spines with classic mushroom heads, markers of matured spines, was increased. Finally, the proportion of branched spines was significantly decreased in *Pyk2*-KO mice. Taken together, we suggest that Pyk2 plays an important role in regulating cytoskeleton remodeling and spine dynamics.

The clustered Pcdh proteins are essential for self-avoidance and tiling of neuronal processes as well as proper assembly of neuronal connectivity [6,7,14,15,43]. These Pcdhs negatively regulate cell-adhesion kinases Pyk2 and Fak through the cytoplasmic domains [5–7]. The present study found a novel fear memory suppression function of Pyk2 through the kinase-independent scaffolding pathway to IEGs. Interestingly, Pyk2 suppresses contextual fear memory but not the learning process. Recent studies revealed distinct hippocampal pathways for contextual fear learning and memory [44,45]. Future studies will be focused on the identification of hippocampal memory engram cells for this Pyk2-related contextual fear circuitry.

## Materials and methods

### Establishment of *Pyk2* gene modified mice

Animals were maintained at 23°C in a 12-h (7:00-19:00) light and 12-h (19:00-7:00) dark schedule. All experimental procedures were performed in accordance with Institutional Animal Care and Use Committee guidelines of Shanghai Jiao Tong University. All mice were deeply anesthetized before sacrifice. Mouse lines deficient for Pyk2 protein were generated using CRISPR/Cas9 system [11,46]. Briefly, we first selected an sgRNA-target sequence at the first exon of the *Pyk2* locus, and then a T7 promoter containing-sgRNA PCR product was amplified from pLKO.1-sgRNA plasmid using oligos, TAATACGACT CACTATAGGG GGCACTTTAC GCCGGCCTGA GTTTTAGAGC TAGAAATAG and AAAAGCACCG ACTCGGTGCC, as forward and reverse primers, respectively. The resultant product was gel-purified and used as a template for *in-vitro* transcription using MEGA short script T7 kit (Life Technology). Cas9 mRNA was transcribed *in vitro* from linearized pcDNA3.1-Cas9 plasmid using mMACHINE T7 ULTRA kit (Life Technology). Both Cas9 mRNA and sgRNAs were purified using MEGA clear kit (Life Technology) and mixed in RNase-free TE buffer at the concentration of 100 ng/µl. After equilibration for 30 minutes, 15-25 injected embryos were transferred into the Fallopian tube of pseudopregnant ICR females for generating chimeric mice. Germline transmission and genotyping were confirmed by PCR using allele-specific primers (Forward: GGATGTGGCA TGTGGCTTGC AAGAG; Reverse: TACCTGGATC TCTGTCTGCA CTGTG) under the following conditions: 94°C for 2 min; (94°C for 30 s, 59°C for 30 s, 72°C for 30 s) for 35 cycles; 72°C for 3 min; hold at 4°C. PCR products were then gel-purified, and digested with the Bsll enzyme. Digested DNA was separated on an Ethidium bromide stained Agarose gel. For identification of mutations, purified PCR products were used for TA cloning and subsequently for Sanger sequencing. The method for establishing Pyk2^Y402F^ mutant mice was similar to that of *Pyk2-* KO mice with sgRNA-target sequence designed at the 13^th^ exon of the *Pyk2* locus using oligos, TAATACGACT CACTATAGGG TACAGAGTCA GACATCTATG GTTTTAGAGC TAGAAATAG and AAAAGCACCG ACTCGGTGCC, as forward and reverse primers, respectively. Furthermore, single-stranded oligonucleotides (S2 Fig) were used as a donor DNA with a mutation at Y402 residue and a nonsense mutation at the PAM sequence for co-injection into the fertilized eggs together with Cas9 mRNA and sgRNA. The germline transmitted mutants were identified by PCR using allele-specific primer pairs (Forward: ACTGTGTGGC TTCCTTGAAT CCTGG; Reverse: TCTCCTGTGG TGTCCCATGA ATAC) under the following conditions: 94°C for 2 min; (94°C for 30 s, 59°C for 30 s, 72°C for 30 s) for 35 cycles; 72°C for 3 min; hold at 4°C. PCR products were then-gel purified and confirmed by Sanger sequencing.

### Behavior tests

All behavioral experiments were performed blind to genotypes of mice, and independent experimenters analyzed the data. For contextual and auditory fearing conditioning, tests were performed as previous published [26,28] and the protocol was illustrated in Fig 2A. Briefly, mice were put into the training chamber for 12 minutes of adaptive training on Day 1. Fear training was performed on Day 2 as the following: Mice were allowed to acclimate to the chamber for 4 min prior to six cycles of consecutive training, each consisting of a 20s baseline, an auditory tone, 18s of trace interval, 2s of foot shock, and followed by a 40s inter-trial interval (ITI). On Day 3, mice were put back to the original training chamber for 3 min to assess contextual fear conditioning, after which they were returned to their home cage for 3 min. For trace fear conditioning testing, mice were then placed in a novel chamber different from the training chamber for 3 min as an index of novel contextual fear responses. This was followed by an auditory trace response testing. Percentage of time spent freezing was recorded using automated Video Freeze software.

For Morris water maze tests, experiments were performed using the protocol previously published [28]. Briefly, mice were tested in a pool of 120 cm in diameter surrounded by different visual cues. A 10 cm-diameter circular platform was used as the goal platform. Room and water temperature were maintained at 22–23°C. Testing includes three stages of training, acquisition, and reversal. Mice were handled 5 minutes for consecutive 3 days by experimenters for adaptive training followed by 2 days of training, in which testing was conducted 4 trials each day and the platform was put visibly at different quadrants for each trial. In the acquisition sessions, the platform was not visible and placed in a fixed location for 4 days. Mice were tested in four trials per day, with a 10-min interval between each trial. Then the platform was put into the opposite quadrant for reversal testing for another 5 days. For each trial per day, mice were placed into the pool at start positions distributed in different quadrants, and the latency to the platform was automatically recorded using the Clever system software (TopScan3.0).

### RNA sequencing and data analyses

Hippocampus collected from wild-type and gene-modified mice were used for total RNA extraction using TRIzol reagent according to the manufacturer’s protocol (Life Technology). The concentration of the RNA was measured by using NanoDrop 2000. At least 200 ng of total RNA was used for preparing RNA libraries before deep sequencing according to the manufacturer’s instructions (Illumina, USA). The cDNA libraries were sequenced on Illumina HiSeq 4000 instrument with 50-base single reads. Raw sequencing data was cleaned and mapped to the mouse reference genome using Bowtie2. Relative abundance of transcripts was measured by Fragments Per Kilobase of exon per Million mapped fragments (FPKM). Differentially-expressed mRNAs between gene-modified and wild-type mice were identified using the Cuffdiff module, and genes with >2-fold changes or <0.5-fold changes and FDR<0.001 were selected.

### Golgi staining and 3D reconstruction of CA1 pyramidal neurons

Golgi staining experiments were performed as previously described [7]. Briefly, *Pyk2-*KO and wild-type littermates were anesthetized and sacrificed. Brains were collected as quickly as possible and put into the impregnation solution for 14 days. The impregnation solution was then replaced by the Solution C for another 3 days. Brain was then cut into 150 µm sections using a Vibratome (Leica VT1200S). After dehydration, sections were cleared in xylene before mounting using Permount (Fisher Chemical™). Images were collected with Nikon confocal microscopy (Nikon A1) under a 20x objective for dendritic analyses using the Neuromantic software (http://www.reading.ac.uk/neuromantic). Each CA1 pyramidal neuron was 3D-reconstructed by tracing the dendritic paths of the Z-stacking pictures. The number of branch points (bifurcations) and of segments between branch points were analyzed automatically by the software. For spine analyses, high-resolution images were collected under a 60x oil objective with a 3X digital zooming factor. The reconstructed Z-stacking pictures were imported into the Image J (NIH) for analyzing the spine morphology. The experimenter and analyzer were blinded to genotypes during data analyses.

### Immunohistochemistry

Brains removed from embryonic 18.5, postnatal day 0, and adult mice were fixed in 4% PFA overnight at 4°C before cutting into 50 µm sections with a Vibratome. After permeating with 0.3% Triton-X100 and blocking with 3% BSA, sections were incubated with rabbit anti-Pyk2 antibody (Abcam) at 4°C overnight. Signals were detected with Alexa 488 Fluor-conjugated secondary antibodies. Cell nuclei was visualized with DAPI.

### Hippocampal neuron culture and time-lapse analyses

Hippocampus was collected from E18.5 embryos in Hanks’ Balanced Salt Solution with 0.5% glucose, 10 mM Hepes, and 100 mg/ml penicillin/streptomycin. Tissues were digested with 0.25% trypsin for 15 min at 37°C. After stopping the reaction with trypsin inhibitor (0.5 mg/ml) for 3 min at room temperature, tissues were gently triturated in the plating medium (MEM medium, 10% FBS, 1 mM glutamine, 10 mM Hepes, 50 mg/ml penicillin/streptomycin). Cell viability and density were determined using 0.4% trypan blue and a hemocytometer. Cells were plated at the density of 500 cells/mm_2_ onto glass coverslips with a uniform application of poly-D-lysine and laminin (Corning BioCoat). Cells were incubated with 5% CO_2_ at 37°C. After 3–4h, the plating medium was replaced with a serum-free culture medium (Neurobasal medium, 2% B27, 0.5 mM glutamine, 50 mg/ml penicillin/streptomycin supplemented with 25 mM glutamate acid), and thereafter half of the medium (W/O glutamate) was replaced every 3 days. On Day 3, 2 mM cytosine arabinoside (Sigma) was added to inhibit the proliferation of glia. Neurons were transfected with pCAG-EGFP plasmids at 12 DIV using a Lipofectamine 2000 (Invitrogen) and cultured until 14 DIV before fixing for analyses. For immunofluorescent staining, cultured primary hippocampal neurons were washed with 1x PBS, fixed in 4% PFA for 20 min at room temperature, washed again and permeabilized with 0.3% Triton X-100 for 20 min. After blocking with 3% BSA, cells were incubated with rabbit anti-GFP antibody (Invitrogen) at 4°C overnight. Signals were detected with Alexa 488 Fluor-conjugated secondary antibodies. Cell nuclei were visualized with DAPI. For spine analyses, high-resolution images were collected under a 60x oil objective with a 3X digital zooming factor. For time-lapse analyses, transfected neuronal cells were maintained at physiological conditions with a stage-top microscope incubator, and Z-stack images were acquired on an inverted laser-scanning confocal microscope. Pictures were captured every 1 min for up to 1h with a 60x, 1.4 numerical aperture objective.

### Protein extraction for western blotting

Hippocampus from adult mice were homogenized with the RIPA lysis buffer (50 mM Tris–HCl, pH 7.5, 150 mM NaCl, 1% NP40, 0.25% sodium deoxycholate, 10 mM NaF, 10 mM Na_3_VO_4_, 1 mM phenylmethanesulfonyl fluoride (PMSF), and protease inhibitors) at 20 µl/mg ratio. The homogenate was centrifuged at 12,000*g* for 20 min and the resulting supernatant was collected and concentrated with the BCA method. Rabbit anti-Pyk2 antibody (Abcam), mouse anti-β-actin antibody (Proteintech), IRDye680-conjugated goat anti-rabbit and IRDye800-conjugated goat anti-mouse secondary antibodies (both from Biosciences) were used for Western blot. Signals were scanned on the Odyssey system (Li-Cor).

### Statistics

Data were analyzed by GraphPad Prism software using two-way ANOVA with repeated measures. Bonferroni post tests were used for pairwise comparison. Data for contextual fear conditioning and dendritic morphology analyses between wild-type and Pyk2 mutant mice were analyzed using a two-tailed, unpaired Student’s *t* test. Data were represented as mean ± SEM. *p < 0.05, **p < 0.01, ***p < 0.001.

## Acknowledgments

We thank Wuming Wang for the initial work on the project, Prof. Weidong Li (SJTU) for allowing us to use his fear-conditioning testing equipment, Dr. Wenjie Bian (ION) for advice on the Golgi staining, and Prof. Aaron Hsueh (Stanford) for linguistic help on the manuscript.

## Author Contributions

**Mice generation:** LS YXZ.

**Study design:** LS JZ YXZ QW.

**Primary culture and biochemistry experiment:** LS YXZ.

**Animal behavior tests:** LS JZ LLJ.

**Golgi staining and picture collection:** LS JZ.

**Data analyses:** LS YXZ JZ.

**Study supervision:** YPK QW.

**Manuscript preparation:** LS QW.

